# Efficient genomic prediction at reduced training size and moderate marker density in an expanded aus-NAM population of rice

**DOI:** 10.64898/2026.04.28.721500

**Authors:** Justine K. Kitony, Vincent P. Reyes, Hidehiko Sunohara, Mikako Tasaki, Masanori Yamasaki, Jun-ichi Mori, Akihisa Shimazu, Shunsaku Nishiuchi, Todd P. Michael, Kazuyuki Doi

## Abstract

Genomic selection (GS) can accelerate genetic gain in crops, but its effectiveness depends on training population design and marker density. Nested association mapping (NAM) populations provide a structured framework that captures broad allelic diversity within a controlled genetic background. Here, we evaluated genomic prediction (GP) and genome-wide association study (GWAS) performance in an expanded aus-NAM population of rice comprising 1,818 recombinant inbred lines across 14 families and 11 agronomic traits, using genotyping-by-sequencing (GBS) markers and projected whole-genome sequence variants. Prediction accuracy plateaued at moderate marker densities (∼20k SNPs) and with training populations of ∼500 lines (∼40–60% of the available pool), with trait heritability emerging as the strongest determinant of predictive performance rather than model choice or marker density. In contrast, GWAS resolution continued to improve with increasing marker density, enabling detection of additional loci, including a chromosome 12 locus associated with heading date, while consistently recovering well-characterized genes such as *EARLY HEADING DATE 1 (Ehd1)* and *SEMIDWARF 1 (SD1)*. These contrasting patterns indicate that GP reaches near-optimal performance once genome-wide variation is adequately represented, whereas GWAS benefits from higher marker density through improved locus resolution. The present study establishes a benchmark for implementing breeding programs involving japonica/indica crosses using GP in a single environment.

## Background

Meeting rising global food demand requires advances in breeding methodologies, as current rates of genetic gain remain insufficient to close the gap between crop production and projected demand (Khush 2013; Duan et al. 2024; Voss-Fels et al. 2019). Genomic selection (GS), also referred to as genomic prediction (GP), has emerged as a promising strategy to accelerate breeding by predicting the genetic merit of individuals using genome-wide markers (Pérez and de los Campos 2014; Crossa et al. 2017). Although GS has demonstrated success in specific contexts, including drought-tolerant maize and improved parental selection in oil palm (Gaffney et al. 2015; Aleri et al. 2026; Beyene et al. 2015; Kwong et al. 2017), its broader and consistent implementation remains variable and often limited by factors such as population design, training set composition, and genotype-by-environment interactions (Bartholomé et al. 2022; Rice and Lipka 2021).

Rice (*Oryza sativa* L.), a staple crop for more than half of the world’s population, represents a critical target for improving breeding efficiency under diverse and changing environments (Fukagawa and Ziska 2019; Duan et al. 2024). Many agronomic traits in rice, including heading date, plant height, yield components, amylose content, and biomass, are complex quantitative traits controlled by numerous loci and strongly influenced by environmental conditions (Nguyen et al. 2023; Lu et al. 2025). Conventional breeding relies on extensive multi-environment phenotyping, making selection time-consuming and costly (Guimarães 2009). The effectiveness of GP in addressing these challenges depends on factors such as training population design, marker density, genetic architecture, and genotype-by-environment interactions (Crossa et al. 2017; Norman et al. 2018). Structured populations, particularly nested association mapping (NAM), address this need by combining controlled genetic structure with broad allelic diversity. In our previous study, we developed an aus-NAM population consisting of multiple recombinant inbred line families derived from crosses between diverse aus donors and the common parent T65 (japonica) (Kitony et al. 2021; Reyes et al. 2022; Kitony 2023). The aus group was selected as they are considered a reservoir of adaptive alleles, including photoperiod insensitivity, drought tolerance, and disease resistance (Garris et al. 2005; Takehisa et al. 2009; Travis et al. 2015; Norton et al. 2018). Building on this framework, the population was expanded from seven to fourteen families, increasing allelic diversity and population size and providing an enhanced resource for dissecting complex traits in rice.

Despite progress in rice GP, few studies have systematically disentangled the contributions of marker density, training population size, and model choice to prediction accuracy within the same structured population and trait set, information that is essential for designing cost-effective implementation strategies. Here, we address this gap using the expanded aus-NAM population across 11 agronomic traits and SNPs ranging from 1k to 56k markers, identifying the practical thresholds at which prediction accuracy saturates and showing that these thresholds differ fundamentally from those that govern GWAS resolution. We further show that the same population and marker data simultaneously support improved GWAS resolution — and that the divergent response of prediction and mapping to increasing marker density has a practical consequence: a two-tiered genotyping strategy that makes full use of the NAM framework without requiring all lines to be genotyped at maximum density.

## Materials and Methods

### Plant materials and phenotyping

The aus-NAM population comprises 2,264 recombinant inbred lines (RILs; Supplemental Data Table 1) derived from 14 biparental families sharing the elite japonica cultivar Taichung 65 (T65) as a common parent.

Phenotypic data were collected in 2018 at Togo Field, Nagoya University, Aichi, Japan (35°06’36.5“N, 137°05’06.3“E). Four seedlings per line per row were grown with a spacing of 20 cm between the hills and 30 cm between rows. Standard agronomic management was followed during the experiment, except that no fertilizer was applied. A total of 11 agronomic traits: days to heading (DTH), culm length (CL), panicle length (PL), panicle rachis length (PRL), panicle number (PN), panicle weight (PW), shoot weight (SW), number of primary branches (NPB), number of spikelets (NSSP), and seed setting rate (SSR). Biomass (BM) was calculated as PW + SW. Trait-wise analyses excluded lines with missing phenotypes. Pearson correlations among traits were computed using complete observations.

### Genotyping and SNP processing

Genomic DNA was extracted from approximately 5 cm of leaf tissue per line using a modified Dellaporta method (Dellaporta et al. 1983). Samples were oven-dried at 53 °C overnight before extractions. DNA quality and quantity were assessed by agarose gel electrophoresis (0.6% TBE) and the Quantiflour dsDNA system (Promega, USA).

Genotyping-by-sequencing (GBS) libraries were prepared following established protocols (Poland et al. 2012; Furuta et al. 2017). Briefly, 200 ng of genomic DNA per sample was double-digested with KpnI and MspI enzymes (New England Biolabs, USA), ligated to barcode adapters, multiplexed, purified (QIAquick PCR purification kit, Qiagen), PCR-amplified, and sequenced on an Illumina platform (Poland et al. 2012; Furuta et al. 2017; Reyes et al. 2022).

Raw sequence reads were processed using the TASSEL-GBS pipeline v5 (Glaubitz et al. 2014) and aligned to the Oryza sativa Nipponbare reference genome (IRGSP-1.0; (Kawahara et al. 2013)). SNPs were filtered using the following criteria: minor allele frequency (MAF) ≥ 0.02, minimum locus coverage ≥ 0.3, missing rate ≤ 0.5, and heterozygosity ≤ 0.125. Only SNPs polymorphic between parental lines and monomorphic within each parent were retained. Missing genotypes were imputed using the FSFHap algorithm (Swarts et al. 2014) implemented on TASSEL-GBS v5 (Glaubitz et al. 2014).

After filtering, a total of 1,818 recombinant inbred lines (RILs) across 14 families were retained, with SNP counts per family ranging from 2,522 to 5,019 (Table 1). Raw SNP counts ranged from 9,871 to 19,434, reflecting variation in sequencing depth and parental diversity.

**Table 1.** Summary of *aus-NAM* population and genotype dataset.

| Family | Donor | Taxa_F5 | Final_taxa | Raw_SNPs | Final_SNPs |
| --- | --- | --- | --- | --- | --- |
| WNAM02 | Kasalath | 114 | 110 | 19434 | 4399 |
| WNAM29 | Kalo Dhan | 135 | 131 | 9871 | 2799 |
| WNAM31 | Shoni | 151 | 147 | 11265 | 3211 |
| WNAM35 | ARC5955 | 157 | 153 | 11546 | 2522 |
| WNAM39 | Badari Dhan | 170 | 166 | 13768 | 3732 |
| WNAM04 | Jena035 | 175 | 171 | 13635 | 3919 |
| WNAM27 | Nepal 8 | 126 | 122 | 12756 | 2980 |
| WNAM28 | Jarjan | 94 | 90 | 12293 | 3700 |
| WNAM30 | Anjana Dhan | 92 | 88 | 10975 | 3359 |
| WNAM32 | Tupa 121 | 157 | 153 | 18098 | 5019 |
| WNAM33 | Surja Mukhi | 135 | 131 | 13756 | 3444 |
| WNAM34 | ARC7291 | 174 | 170 | 11345 | 3276 |
| WNAM36 | Ratul | 134 | 130 | 11033 | 3408 |
| WNAM40 | Nepal 555 | 60 | 56 | 11333 | 3555 |

To obtain high-density SNP information, genomic DNA from the parental lines was used to prepare sequencing libraries with the TruSeq DNA LT Sample Prep Kit (Illumina, San Diego, CA, USA), and paired-end sequencing was performed on the Illumina MiSeq platform using the MiSeq Reagent Kit v3 (600 cycles). After read processing, alignment, and variant filtering, approximately 56k SNPs were identified. To generate a high-density SNP panel for the NAM population, SNP variants identified from parental resequencing were projected onto individual RIL genomes within each biparental family. Projection was performed independently per family to preserve the two-founder haplotype structure inherent to each cross. The procedure followed two steps. First, GBS-derived SNPs were used as anchor markers to define haplotype block boundaries along each RIL genome. Second, for each interval between adjacent anchor markers, if both flanking markers were homozygous and concordant with the same parental allele, all parental WGS-derived SNP variants within that interval were assigned the corresponding parental genotype. Intervals where flanking anchor markers were heterozygous, missing, or discordant — indicating a recombination breakpoint or insufficient marker coverage — were left as missing rather than assigned probabilistically. This procedure yielded a projected panel of approximately 56k SNPs per family, which was subsequently merged across families and used alongside the GBS dataset for GP and GWAS analyses at varying marker densities. To evaluate the effect of marker density, marker sets of approximately 1k, 2k, 5k, 10k, 20k, and 40k SNPs were generated from the 56k panel and are presented as marker density panels.

### SNP heritability

Marker-based heritability (h2) was estimated using the 10k genotype panel employed in GP. Genomic relationship matrix (GRM) was constructed using the rrBLUP v.4.6.3 framework (Endelman 2011). Variance components were estimated using a mixed model with family fitted as a fixed effect and additive genomic effects fitted as a random effect. SNP heritability was calculated as h²=Vg/(Vg+Ve), where Vg is the additive genetic variance captured by markers, and Ve is the residual variance.

### Population structure

Genotypes were encoded as −1, 0, and 1, for homozygous reference, heterozygous, and homozygous alternative states, respectively, with missing genotypes retained as NA. Principal component analysis (PCA) was performed using pcaMethods v.2.2.0 (Stacklies et al. 2007), and the leading components were used to account for population structure.

### Genome-wide association analysis

GWAS results were visualized within a defined genomic window centered on the lead SNP (±2 Mbp) to enable fine-scale interpretation of association signals. SNP significance was plotted as −log₁ ₀ (p-value) against physical position, and linkage disequilibrium (LD; r²) between each SNP and the lead variant was calculated from genotype data and used to color-code markers, highlighting local haplotype structure around the peak.

To capture clusters of association signals beyond single-marker significance, a local score approach based on the Lindley process was applied to transformed statistics (−log₁ ₀ (p-value) − ξ), where ξ was empirically defined (Mercier and Daudin 2001). The cumulative score was computed along genomic coordinates to identify contiguous intervals enriched for association signals, which were used to delineate candidate QTL regions (Bonhomme et al. 2019). SNP-wise contributions to the score were retained to distinguish positive (signal-supporting) and negative (noise-reducing) effects along the region.

Pairwise LD (r²) among SNP loci within each region was further calculated and visualized as a triangular heatmap to characterize local LD block structure and recombination patterns. Concordance among association peaks, local score accumulation, and LD structure was used to refine candidate intervals and support biological interpretation of GWAS signals.

### Genomic prediction

GP was performed using four algorithms: ridge regression best linear unbiased prediction (rrBLUP), reproducing kernel Hilbert space (RKHS), Bayesian B (BayesB), and Bayesian LASSO (BL), implemented in BGLR v1.1.4 (Pérez and de los Campos 2014) and the rrBLUP package (Endelman 2011). Marker genotypes were encoded as −1, 0, and 1, representing homozygous reference, heterozygous, and homozygous alternative alleles, respectively. Prediction accuracy was evaluated using 5-fold cross-validation with 10 repetitions, and calculated as the Pearson correlation between observed and predicted values. Results are reported as mean ± standard error.

### Effect of training population size

To evaluate the effect of training population size, a repeated k-fold cross-validation scheme was implemented. For each trait, individuals were partitioned into five folds, with one fold used as the validation set (∼20%) and the remaining folds forming the training pool (∼80%).

Within each training pool, subsets corresponding to 20%, 40%, 60%, 80%, and 100% of the available individuals were sampled proportionally by family to preserve population structure. Models were trained on each subset and evaluated on the corresponding validation fold.

This procedure was repeated across five folds and ten independent replicates, and prediction accuracy was summarized as the mean ± standard error (SE) across all runs.

### Statistical analyses and visualizations

All analyses were conducted in R v.4.5.2 using packages including rrBLUP, BGLR, pcaMethods, and ggplot2 as well as Python 3.11.7.

## Results

### Phenotypic diversity *of* agronomic traits in the *aus*-NAM population

The aus-NAM population comprises 14 biparental families generated by crossing 14 diverse aus donor lines with T65, creating a structured population that captures broad allelic diversity within a controlled genetic background (Supplemental Fig. 1). Substantial phenotypic variation was observed across all traits, including plant height (Fig. 1a). DTH ranged from 77 to 130 days (mean = 100.2), while CL varied from 32 to 143 cm (mean = 96.3). Yield-related traits also showed wide distributions, including panicle weight (PW; 0.54–67.76 g), shoot weight (SW; 0.18–108.41 g), and biomass (BM; 0.85–139.16 g), indicating broad phenotypic diversity captured within the population (Supplemental Fig. 2 and Supplemental Data Table 1).

**Fig. 1.**
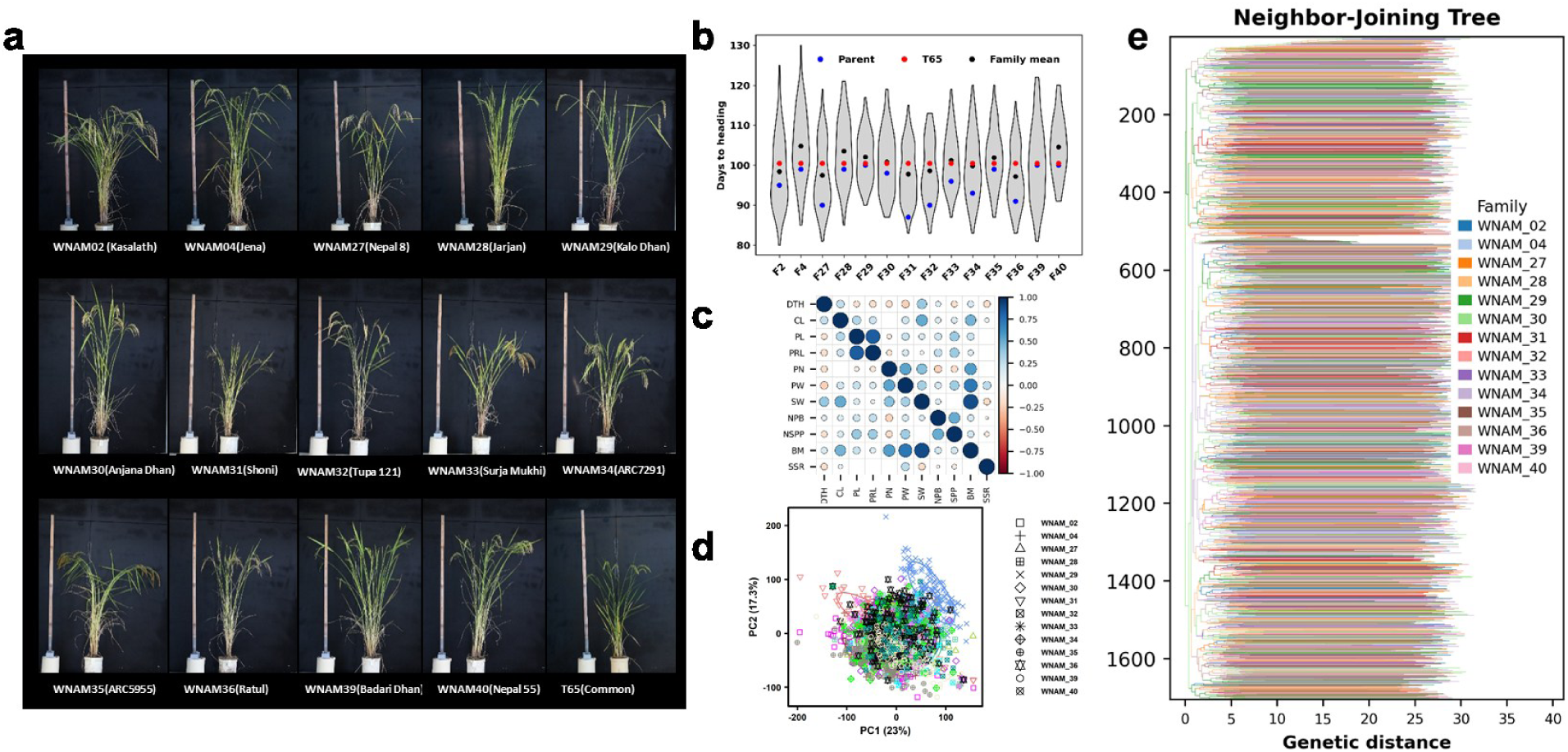
Phenotypic diversity and population structure of the aus-NAM population. (a) Representative images of plant height at maturity for the common parent, T65, and 14 aus donor parents (founders of the NAM population). A ruler is shown for scale. (b) Distribution of days to heading (DTH) across 14 NAM families. Each violin represents within-family variation among recombinant inbred lines (RILs). Black dots indicate family means, blue dots represent donor parents, and the red dot denotes the common parent (T65). (c) Correlation matrix of agronomic traits measured in 2018. Circle size and color indicate the magnitude and direction of Pearson correlations (blue = positive, red = negative). Non-significant correlations (P > 0.05) are omitted. (d) Principal component analysis (PCA) of the aus-NAM population. Points represent individual RILs colored by family, and ellipses indicate group dispersion. (e) Neighbor-Joining tree of the aus-NAM population (14 families) based on genome-wide SNP data. Branches are colored by family. Clear clustering by family is not strictly resolved.

Despite the shared genetic background, some transgressive segregation was observed across multiple families. For example, in several families (WNAM04, WNAM28, WNAM29, WNAM30, WNAM33, WNAM35, and WNAM40), family mean DTH exceeded both parental values, demonstrating the recombination of alleles generating novel phenotypes beyond the parental range (Fig. 1b). Notably, the common parent T65 consistently flowered earlier than all aus donor parents, highlighting its role in shaping phenological variation within the population.

Trait correlation analysis revealed consistent relationships among traits (Fig. 1c). DTH was positively correlated with SW, which in turn was strongly correlated with BM. In contrast, DTH was negatively correlated with PN and PW. Principal component analysis further indicated limited population structure, with PC1 and PC2 explaining 23% and 17.3% of the genetic variation, respectively (Fig. 1d). Higher-order components (PC3–PC5) also contributed appreciably (17.0%, 15.6%, and 14.0%), reflecting the multi-parent design of the NAM population (Supplemental Fig. 3). Most lines clustered according to family background but with substantial overlap, confirming that the population captures broad diversity within a controlled genetic framework. A subset of lines from WNAM29 showed partial separation from the main cluster.

Consistent with PCA, the neighbor-joining tree revealed clear clustering of lines within families while maintaining overall connectivity across the population due to the shared common parent T65 (Fig. 1e).

### Prediction accuracy across marker densities

Prediction accuracy varied across traits, marker densities, and models (Fig. 2). Across all analyses, DTH showed the highest prediction accuracy, ranging from 0.54 to 0.64 depending on marker density and model (e.g., BayesB: 0.58 at 5k to 0.64 at 40k; rrBLUP: 0.57 at 5k to 0.61 at 20k). Other traits exhibited moderate accuracies, including CL (0.48–0.58), PL (0.46–0.51), and NSPP (0.40–0.49), whereas traits such as SSR consistently showed low prediction accuracy (0.14–0.19) across all marker densities and models.

**Fig. 2.**
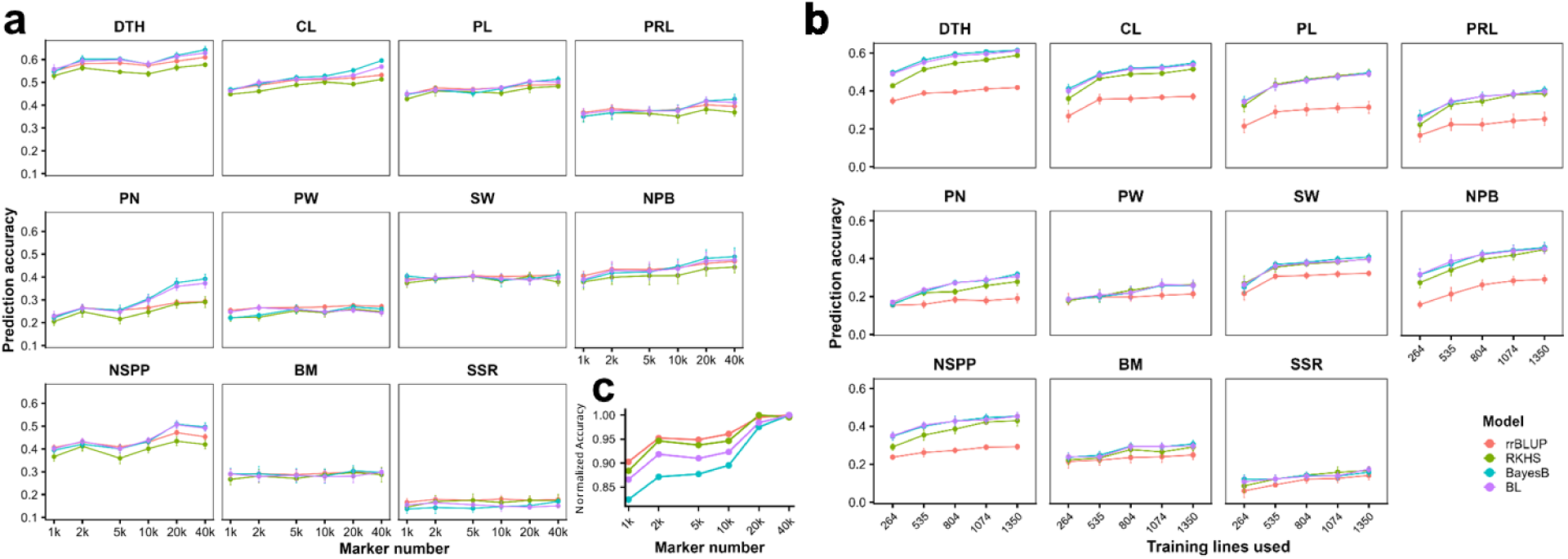
Prediction accuracy plateaus at moderate marker density and reduced training population size in an *aus-*NAM population. (a) Prediction accuracy (y-axis) across marker densities (1k–40k SNPs), faceted by trait, for four models (rrBLUP, BayesB, BL, RKHS) across 11 agronomic traits: days to heading (DTH), culm length (CL), panicle length (PL), panicle rachis length (PRL), panicle number (PN), panicle weight (PW), shoot weight (SW), primary branches (NPB), spikelets (NSSP), and seed setting rate (SSR); biomass (BM) was calculated as PW + SW. (b) Prediction accuracy as a function of training population size (number of training lines). Error bars indicate ±SE across cross-validation replicates. (c) Normalized mean prediction accuracy averaged across all traits as a function of marker density (1k–40k SNPs).

Increasing marker density from 5k to 10k SNPs resulted in the largest gain in prediction accuracy across most traits. For example, DTH improved from 0.57 to 0.60 (rrBLUP) and from 0.58 to 0.62 (BayesB), while CL increased from ∼0.49 to ∼0.53. Further increases in marker density (20k–40k) resulted in smaller changes in prediction accuracy. For DTH, prediction accuracy ranged from 0.60 to 0.63 across higher marker densities, with no consistent increase beyond 10k–20k markers and slight reductions in some models (e.g., rrBLUP: 0.61 at 20k vs 0.59 at 40k). Similar patterns were observed for PRL, SW, and BM.

Trait-specific differences in response to marker density were observed. DTH reached similar levels of prediction accuracy across marker densities above 10k SNPs. CL, PL, and NSPP showed gradual increases in prediction accuracy with increasing marker density, whereas SSR and PW showed limited changes in prediction accuracy across marker densities.

Differences among GP models were generally modest relative to trait effects. Bayesian models (BayesB and BL) showed slightly higher prediction accuracy at higher marker densities (e.g., BayesB reaching a mean accuracy of ∼0.42 at 40k SNPs), whereas rrBLUP showed relatively stable performance across marker densities. RKHS showed variable performance across traits and marker densities without consistent differences relative to other models.

Heritability varied across traits (Fig. 3i), with high estimates for DTH (h² = 0.76) and CL (h² = 0.59), moderate estimates for PL (h² = 0.52), NSPP (h² = 0.50), NPB (h² = 0.56), and SW (h² = 0.37), and lower values for PRL (h² = 0.40) and PN (h² = 0.33). Biomass and yield-related traits, including PW (h² = 0.19), BM (h² = 0.20), and SSR (h² = 0.10), showed relatively low heritability.

**Fig. 3.**
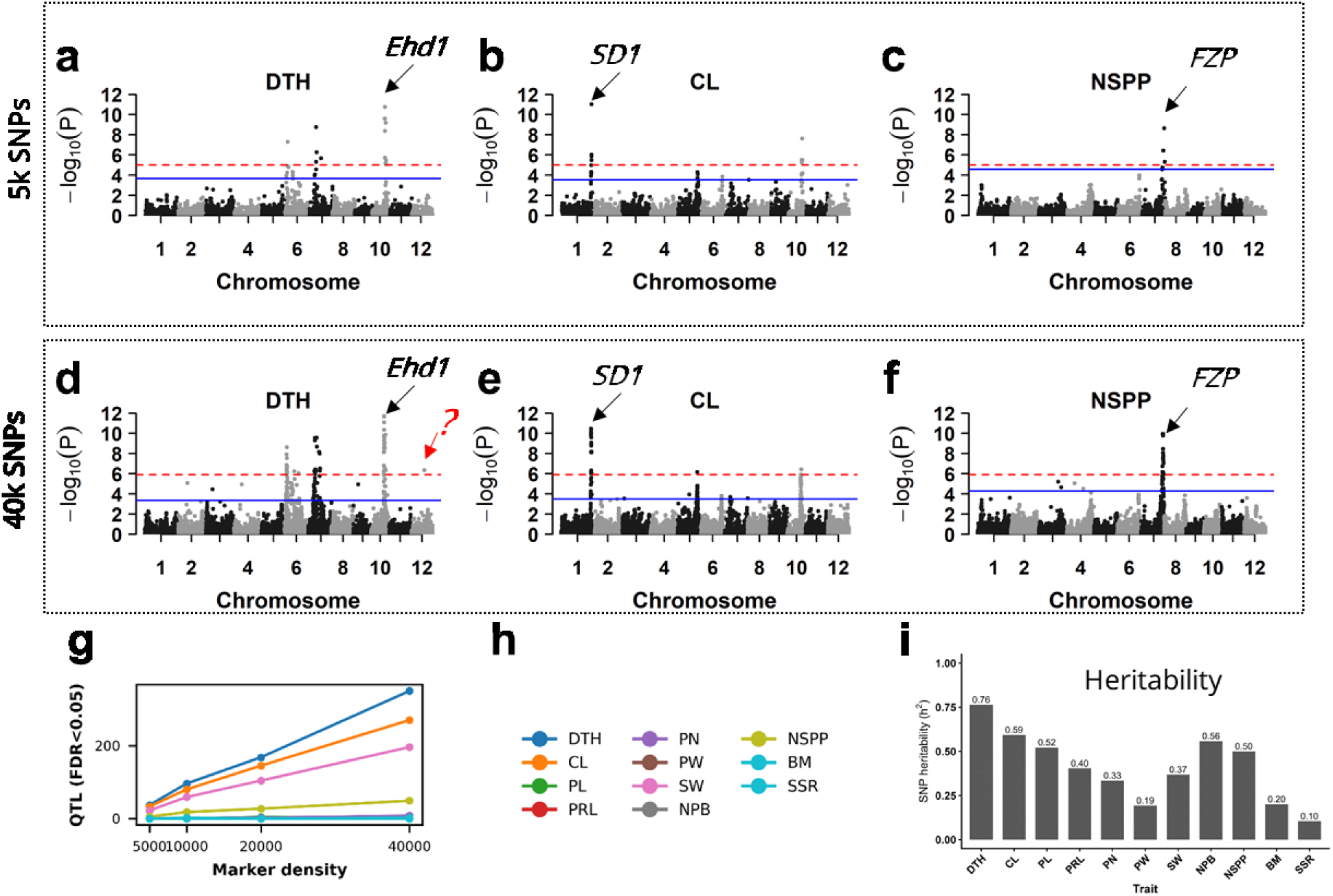
Increased marker density enhances GWAS resolution and QTL detection. (a–c) Manhattan plots for DTH (a), CL (b), and NSPP (c) generated using 5k SNPs. (d–f) Corresponding Manhattan plots using 40k SNPs. Horizontal lines indicate significance thresholds: Bonferroni (red dashed) and FDR (blue). (g) Number of detected QTL (FDR < 0.05) across marker densities (5k–40k SNPs). (h) Legend for (g) and (h). (i) SNP-based heritability (h²) estimates for 11 agronomic traits. Heritability was calculated using marker-based genomic relationships, and values are shown above each bar.

### Prediction accuracy across training population sizes

GP accuracy increased with training population size across all traits and models (Fig. 4b). Notably, using only 20–40% of the training pool (∼260–540 lines) still achieved moderate to high prediction accuracies for several traits. For example, DTH showed accuracies of ∼0.49–0.50 at 20% training size, increasing to ∼0.61 at full training. Similarly, CL achieved accuracies of ∼0.40–0.41 at 20%, rising to ∼0.54 at full training.

**Fig. 4.**
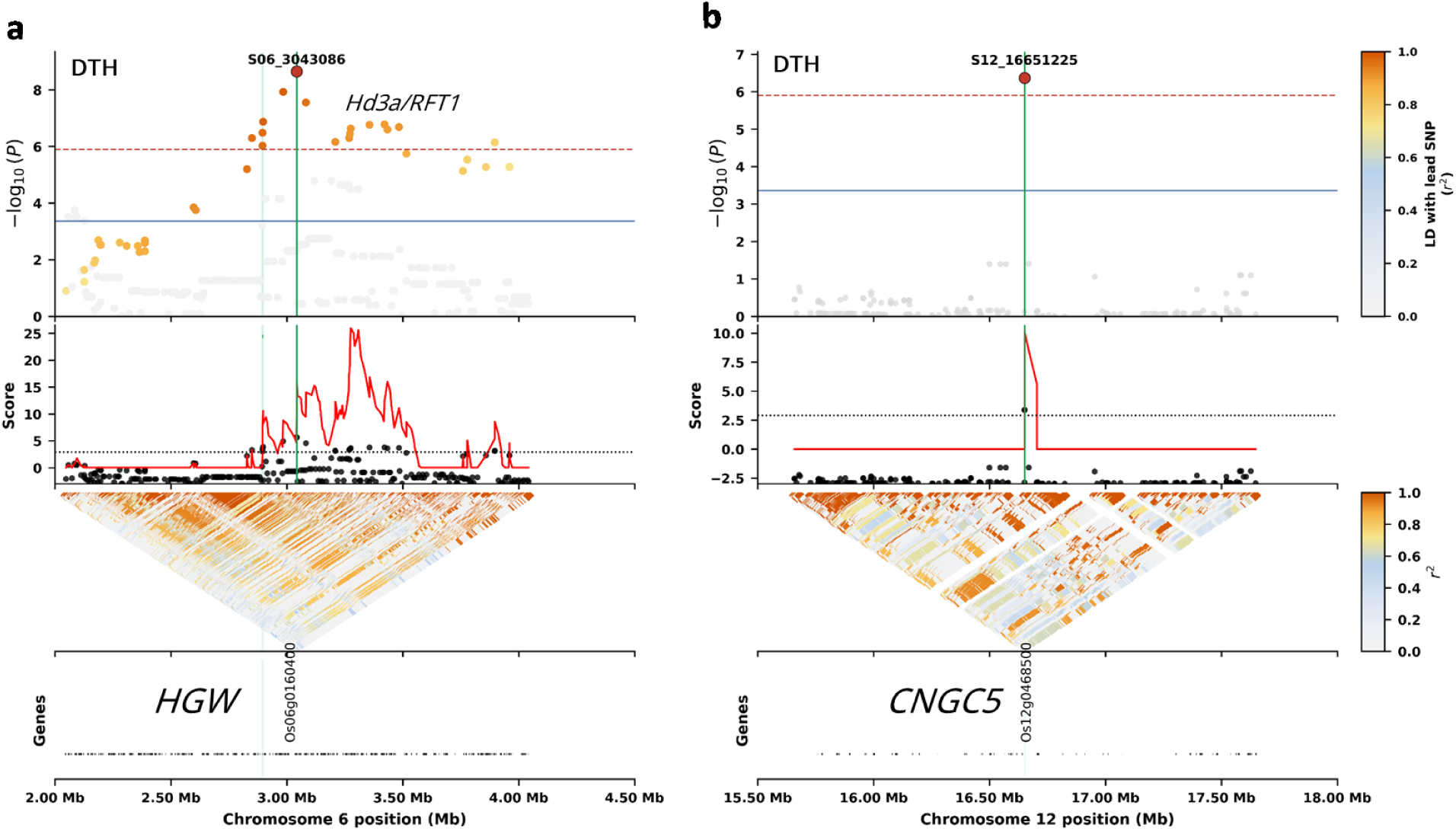
Local GWAS refines candidate regions for heading date. Local GWAS results are shown with SNP significance (−log₁₀(p)) plotted against genomic position and colored by linkage disequilibrium (LD; r²) relative to the lead SNP. The local score profile highlights accumulation of association signals, with the candidate interval indicated by a shaded region. The lower panels show pairwise LD (r²) as triangular heatmaps, revealing local haplotype structure and refinement of association peaks. (a) Chromosome 6 region around S06_3043086 encompassing candidate genes *HGW* (Os06g0160400), *Hd3a*, and *RFT1*; (b) chromosome 12 region around S12_16651225 associated with *OsCNGC5* (Os12g0468500).

Across traits, prediction accuracy generally improved rapidly from 20% to 60% training size and plateaued thereafter, with only marginal gains beyond 80%. This trend was consistent for most traits, including PL, NPB, and NSPP. For instance, PL increased from ∼0.34 at 20% to ∼0.49 at 100%, while NPB improved from ∼0.31 to ∼0.45 across the same range.

Model performance varied across training population sizes. Bayesian models (BayesB and BL) and RKHS achieved higher prediction accuracies than rrBLUP, particularly at smaller training sizes. However, rrBLUP showed similar trends across training population sizes despite lower overall accuracy (Fig. 2b).

### Genome-wide association analysis across marker densities in an aus-NAM population

Increasing marker density increased the number of detected QTL while maintaining the same major signals (Fig. 3). At 5k SNPs, key candidate gene loci were consistently detected, including EARLY HEADING DATE 1 (Ehd1; Os10g0463400), SEMIDWARF 1 (SD1; Os01g0883800), and FRIZZY PANICLE (FZP; Os07g0669500). Increasing density to 40k SNPs identified additional associations, including a DTH locus on chromosome 12 harboring CYCLIC NUCLEOTIDE-GATED CHANNEL 5 (OsCNGC5) (Fig. 4). At this density, the number of significant marker–trait associations (MTAs; FDR < 0.05) varied across traits, with the highest counts observed for DTH (350), CL (270), and SW (196), followed by NSPP (49) and PN (8), while PL (1), NPB (1), BM (2), and PRL, PW, and SSR (0) showed few or no significant associations (Supplemental Data Table 2). Across most traits, the number of significant GWAS signals increased with marker density (Figs. 3g and h).

### Candidate genes underlying QTL identified by GWAS

Local GWAS analyses enabled fine-scale characterization of association signals (Figs. 4 and 5). For DTH, a signal on chromosome 6 (S06_3043086) mapped to a region containing HEADING AND GRAIN WEIGHT (HGW; Os06g0160400) and was proximal to flowering regulators HEADING DATE 3a (Hd3a) and RICE FLOWERING LOCUS T1 (RFT1) (Fig. 4a).

**Fig. 5.**
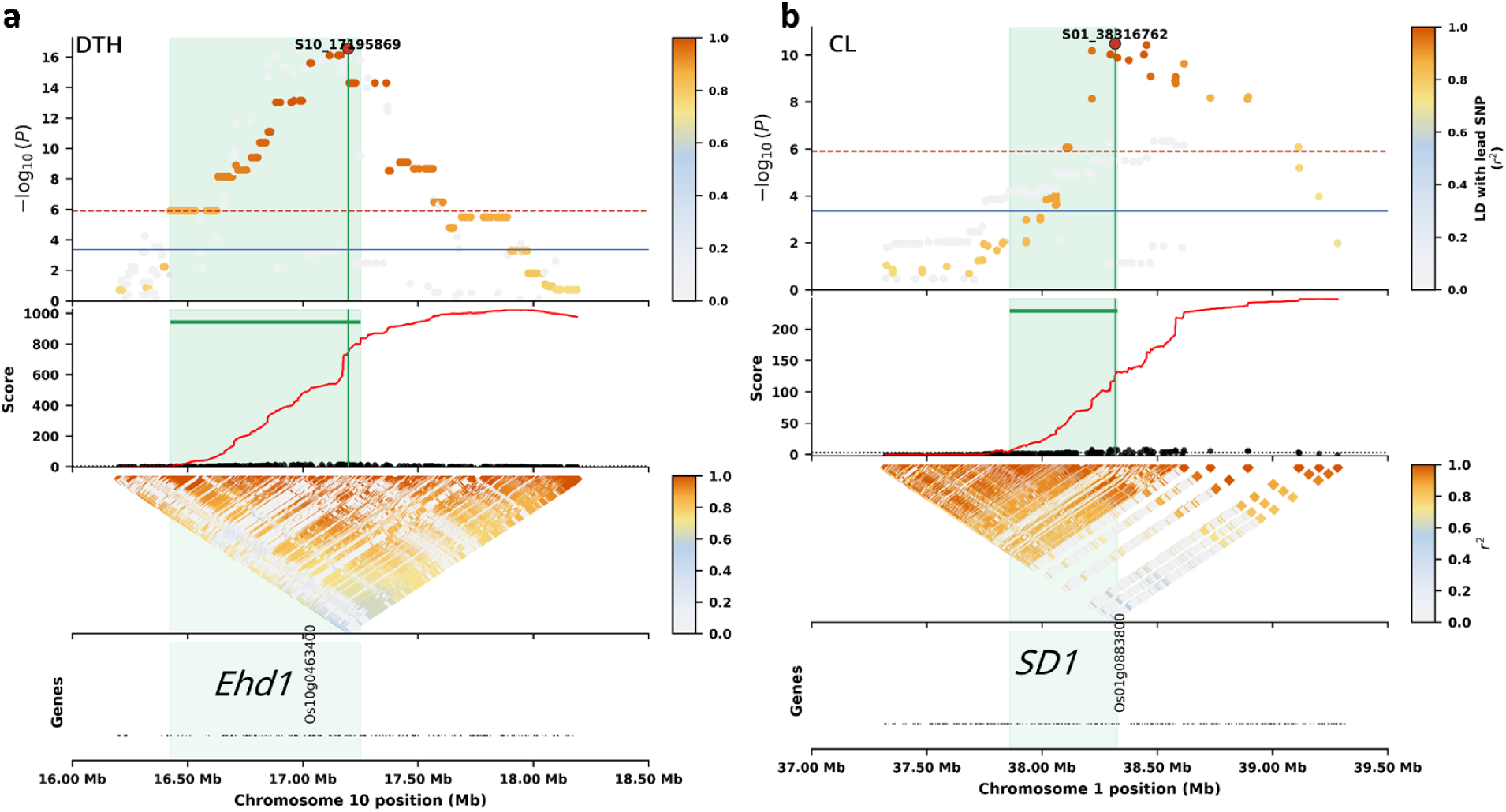
Local GWAS reveals candidate loci underlying key agronomic traits. Local GWAS across genomic position with top panels showing SNP significance (−log₁₀(p); Manhattan plots) colored by linkage disequilibrium (LD; r²) relative to the lead SNP, middle panels showing the local score profile highlighting accumulation of association signal and the refined candidate interval (shaded), and bottom panels showing pairwise LD (r²) as triangular heatmaps, illustrating local haplotype structure and refinement of the association peak. (a) *Ehd1* locus (Os10g0463400) for days to heading (DTH); (b) *SD1* locus (Os01g0883800) for culm length (CL).

On chromosome 7 (S07_10716916), the lead SNP localized near a noncoding transcript, with the closest annotated genes including an ALDEHYDE OXIDASE 3 (OsAAO3; Os07g0282300) (Supplemental Fig. 5a). Consistent with genome-wide results, local GWAS resolved the major locus Ehd1, where dense SNP coverage and local LD structure defined the association peak (Fig. 5a). Similarly, a locus on chromosome 12 (S12_16651225) co-localized with OsCNGC5 (Os12g0468500) and ABERRANT FLOWERING CONTROL 2 (OsAFC2) (Fig. 4b).

For NSPP, a signal on chromosome 7 (S07_27716723) co-localized with FZP (Supplemental Fig. 5b), with nearby genes including CONSTANS-LIKE 13 (OsCOL13) and SPINDLY 6 (SPIN6)

For CL, a signal on chromosome 1 overlapped with SD1 (Fig. 5b).

## Discussion

### The extended aus-NAM population is a suitable platform for studying GP

The aus-NAM population is structured to address a central challenge in rice GP: capturing sufficient allelic diversity for broad applicability while maintaining the controlled genetic background that enables accurate genomic modeling. By deriving all families from crosses with the same recurrent parent, T65, the design generates genome-wide relatedness that supports the estimation of marker effects across families, while the diversity contributed by 14 aus donors ensures that the training population samples a broad range of haplotypes relevant to adaptive traits. This design enables extensive phenotypic variation while minimizing the confounding effects of population structure, supporting both accurate prediction and reliable mapping. The observed clustering by family, with partial overlap among groups, reflects the balance between relatedness and diversity that underpins the utility of NAM populations for quantitative genetic studies (Gage et al. 2020).

The expanded 14-family population delivered on the expectations of the NAM design, with heritability estimates consistent with the known genetic architecture of each trait—high for DTH (h² = 0.76) and CL (h² = 0.59), where major-effect loci such as *Ehd1* and *SD1* are known to segregate in crosses involving aus and japonica parents (Doi et al. 2004; Asano et al. 2011), and lower for yield components such as PW, BM, and SSR (h² = 0.10–0.20), consistent with the high environmental sensitivity documented for these traits in rice (Courtois et al. 2003; Y. Xu et al. 2018). Transgressive segregation was observed across multiple families for DTH (Fig. 1b), confirming that aus donor alleles recombine with T65 haplotypes to generate novel phenotypic combinations beyond the parental range — a well-established property of structured inter-subspecific crosses (Rieseberg et al. 1999). The even distribution of genetic variance across PC1 through PC5 (23.0%, 17.3%, 17.0%, 15.6%, and 14.0%, respectively; Fig. 1d) is characteristic of multi-parent NAM designs, where the shared common parent generates balanced relatedness across families (Yu et al. 2008; Gage et al. 2020). This structure indicates that no single family dominates the population, which is favorable for genome-wide estimation of marker effects.

### GP accuracy is efficiently captured at moderate training population sizes

GP accuracy increased with training population size across all traits and models, with most predictive gain achieved at moderate training sizes of ∼40–60% of the available pool (∼500 lines; Fig. 2b). Beyond this point, accuracy gains were marginal, indicating diminishing returns from additional phenotyping effort. This pattern is consistent with previous findings in structured populations, where training population composition rather than size alone determines prediction performance (Spindel et al. 2015; Yu et al. 2008).

The efficiency of prediction at reduced training sizes reflects the haplotype structure inherent to NAM populations. Because all families share T65 as a common parent, key haplotype blocks and the underlying linkage disequilibrium structure are represented across families even at moderate sampling levels. Once these shared genomic relationships are captured, adding more individuals contributes limited new information to marker effect estimation. The modest, continued improvement observed between 40% and 80% training size likely reflects incremental capture of smaller-effect loci and family-specific recombination events that require greater sampling depth to represent adequately.

Prediction accuracy was evaluated using a repeated k-fold cross-validation framework with proportional family sampling, ensuring that population structure was balanced across training and validation sets. Under this framework, a training set of approximately 500 lines was sufficient to achieve near-maximum prediction accuracy, a finding with direct relevance for programs where phenotyping capacity is the primary constraint on implementing genomic selection (GS).

### Moderate marker density of ∼20k SNPs is sufficient for GP

Prediction accuracy increased from low to moderate marker densities but plateaued at approximately 20k SNPs, beyond which additional markers contributed minimal gains (Fig. 2a). This plateau is consistent with saturation of linkage disequilibrium between markers and causal variants, such that markers beyond this density capture redundant rather than novel genomic information (Yang Xu et al. 2018). Traits with higher heritability reached this plateau earlier and at higher absolute accuracy, whereas traits with lower heritability showed consistently low accuracy regardless of marker density, indicating that the accuracy ceiling for these traits is set by phenotypic quality rather than genotypic resolution. For traits such as DTH and CL, the accuracy gain from 10k to 40k SNPs was less than 0.05 correlation units—a difference unlikely to translate into meaningful improvement in selection decisions and not worth the additional genotyping cost in most breeding programs.

### Heritability of traits determined the GP accuracy more than modeling algorithms

Differences in prediction accuracy across traits were context-dependent (Fig. 2). Although rrBLUP showed lower prediction accuracy under reduced training population sizes, the Bayes based methods (BayesB and BL) and RKHS approaches showed slight advantages in some cases under these conditions. In contrast, when evaluated across marker densities, rrBLUP performed comparably to other models. The lower performance of rrBLUP under reduced training population sizes likely reflects differences in model assumptions. rrBLUP applies uniform shrinkage of marker effects based on an infinitesimal model (Daetwyler et al. 2010), whereas Bayesian models allow variable shrinkage and can better accommodate loci with larger effects (Meuwissen et al. 2001; Habier et al. 2011). Despite this, rrBLUP remains robust and performs comparably to more complex models when genome-wide relationships are effectively captured (Endelman 2011; de Los Campos et al. 2013). Trait-specific patterns further showed that traits with higher heritability exhibited higher prediction accuracy, whereas traits with lower heritability showed consistently lower accuracy across conditions. In practical terms, the choice between rrBLUP and Bayesian models matters most when training data are scarce; once the training population reaches moderate size, the simpler model performs adequately, and the additional computational cost of Bayesian approaches is difficult to justify.

### Statistical power and resolution of GWAS are improved with increasing marker density

GWAS performance improved substantially with increasing marker density (Figs. 3g–h). Higher marker densities increased the number of detected QTL. This reflects enhanced genome-wide coverage and LD tagging, enabling detection of moderate-effect loci that may be missed at lower densities (Gage et al. 2020). This pattern reflects improved genome-wide LD tagging at higher densities, enabling detection of moderate-effect loci that are missed when marker coverage is sparse (Gage et al. 2020). Importantly, major loci including *Ehd1*, *SD1*, and *FZP* were consistently detected across all marker densities, confirming that the underlying genetic signals are robust and that increased density refines resolution rather than altering the fundamental architecture of detected associations.

This contrasts with the behavior of GP, which reaches near-optimal performance at moderate marker densities. The divergence between the two applications — GP plateauing at ∼20k SNPs while GWAS continues to benefit from higher densities — has practical implications for genotyping strategy: moderate-density panels are sufficient for prediction, whereas high-density panels are warranted when the goal is candidate gene resolution (Pang et al. 2020). Compared to large diversity panels such as the japonica-wide GWAS of (Yano et al. 2016), where population size and marker density jointly determine detection power, the NAM design used here achieves consistent detection of major loci within a controlled genetic background, with resolution improving incrementally as marker density increases.

### Known genes underlie major GWAS signals

Local GWAS analyses provided additional resolution by leveraging LD structure and cumulative association signals to refine candidate intervals (Mercier and Daudin 2001). This approach enabled clearer delineation of key loci, including *Ehd1*, *HGW*, and *SD1*, and improved interpretation of association peaks within biologically relevant genomic regions. For example, refinement of the chromosome 12 locus associated with heading date highlighted candidate genes such as *OsCNGC5* and *OsAFC2*, suggesting potential regulatory contributions to flowering time variation (Lee et al. 2022). These results support a genetic architecture in which complex traits are controlled by a combination of major loci and multiple moderate-effect components that can be resolved through increased marker density and localized analysis.

### Application of knowledge obtained in this study to practical breeding

In practical rice breeding, population breeding is gaining popularity over pedigree breeding. The primary difference between these methods lies in the generation at which selection is initiated. GP can be naturally integrated into population breeding, as described by Reyes et al. (2022). The present study establishes a benchmark for implementing breeding programs involving japonica/indica crosses; notably, traits other than DTH were evaluated using a single plant per line, and we characterized the impacts of marker density, population size, and heritability on GP accuracy. Our results indicate that moderate marker densities and optimized training population sizes are sufficient for accurate genomic prediction, whereas higher marker densities are required to enhance GWAS resolution and candidate gene discovery. Although the aus-NAM population utilized only aus varieties as diversity donors and evaluation was done in a single environment, the potential of GP in practical breeding was demonstrated. These findings provide a foundation for integrating GP with complementary approaches, including artificial intelligence, speed breeding, and multi-omics, to accelerate the development of climate-resilient crop varieties (Cobb et al. 2019; Mishra et al. 2018; Crossa et al. 2017; Alemu et al. 2024; Gunundu et al. 2023).

## Declaration

### Ethics approval and consent to participate

Not Applicable

### Consent for publication

Not Applicable

### Availability of data and material

The genotype datasets supporting this study, including genotyping-by-sequencing (GBS)–derived HapMap files for the *aus*-NAM population, parental whole-genome sequence–derived variants, and all supplemental data tables, are publicly available in Figshare at https://doi.org/10.6084/m9.figshare.31931388

### Authors’ contributions

K.D. and J.K.K. conceived the study. K.D., J.K.K., and V.P.R. performed formal analyses. K.D., J.K.K., V.P.R., H.S., M.T., M.Y., J.M., A.S., S.N., and T.P.M. contributed to data interpretation and manuscript revision. All authors read and approved the final manuscript.

## Acknowledgements

We thank the NIAS Genebank for providing founder materials.

## Supplemental Information

Supplemental data are available online.

## Supplemental Figures

**Supplemental Fig. 1.**
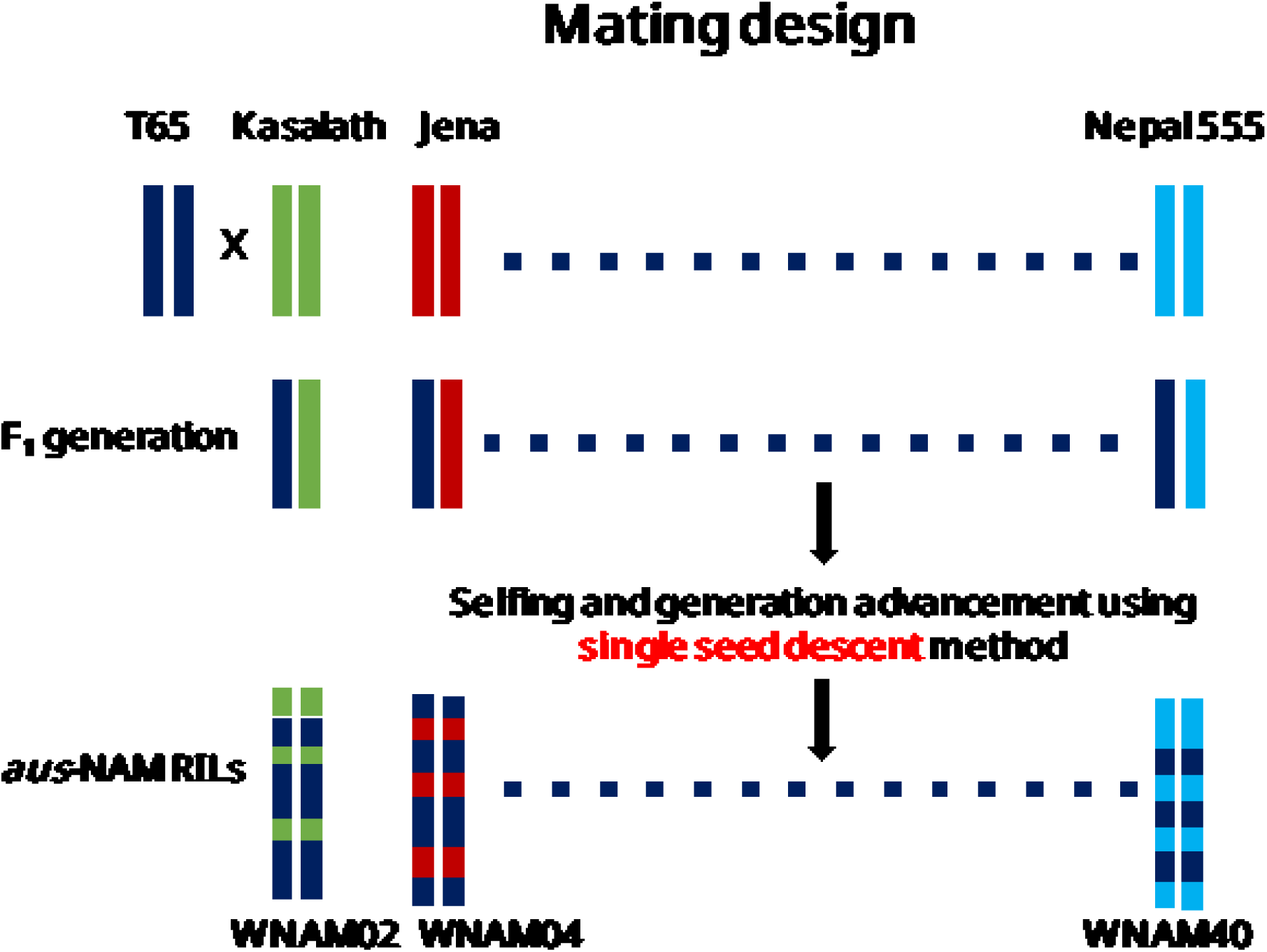
Population design and phenotyping in the aus-NAM population. Breeding scheme of the aus-NAM population. The elite cultivar Taichung (T65) was crossed with 14 diverse aus donor parents to generate bi-parental families, followed by advancement of recombinant inbred lines (RILs) through single seed descent (SSD) to the F5 generation.

**Supplemental Fig. 2.**
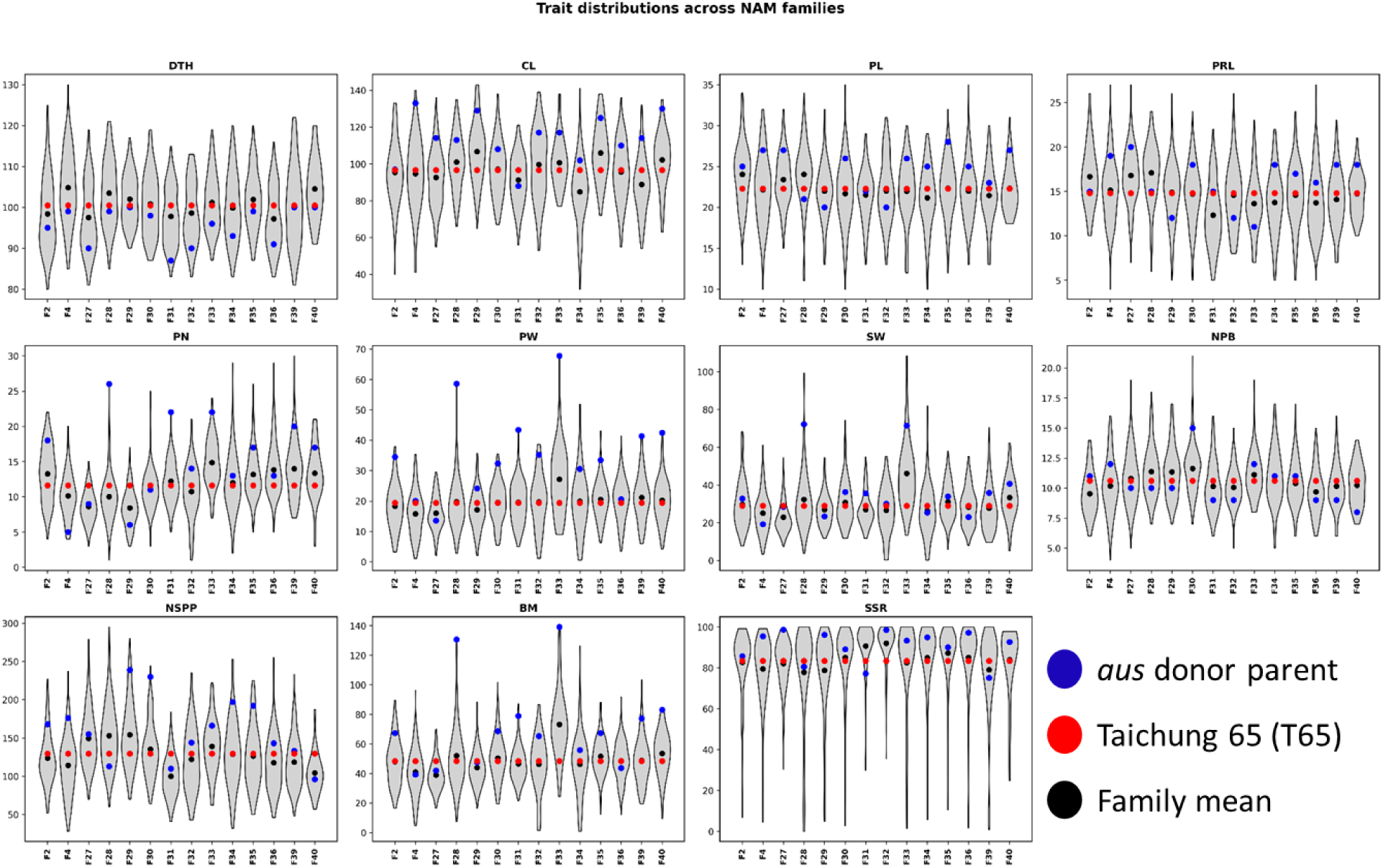
Phenotypic distributions across agronomic traits in the aus-NAM population. Distribution of 11 agronomic traits—days to heading (DTH), culm length (CL), panicle length (PL), panicle rachis length (PRL), panicle number (PN), panicle weight (PW), shoot weight (SW), number of primary branches (NPB), number of spikelets (NSSP), seed setting rate (SSR), and biomass (BM = PW + SW)—across 14 NAM families. Each violin represents within-family variation among recombinant inbred lines (RILs). Black dots indicate family means, blue dots represent aus donor parents, and the red dot denotes the *japonica* common parent (T65). The x-axis represents RIL families ordered from F2 (WNAM02) through F40 (WNAM40).

**Supplemental Fig. 3.**
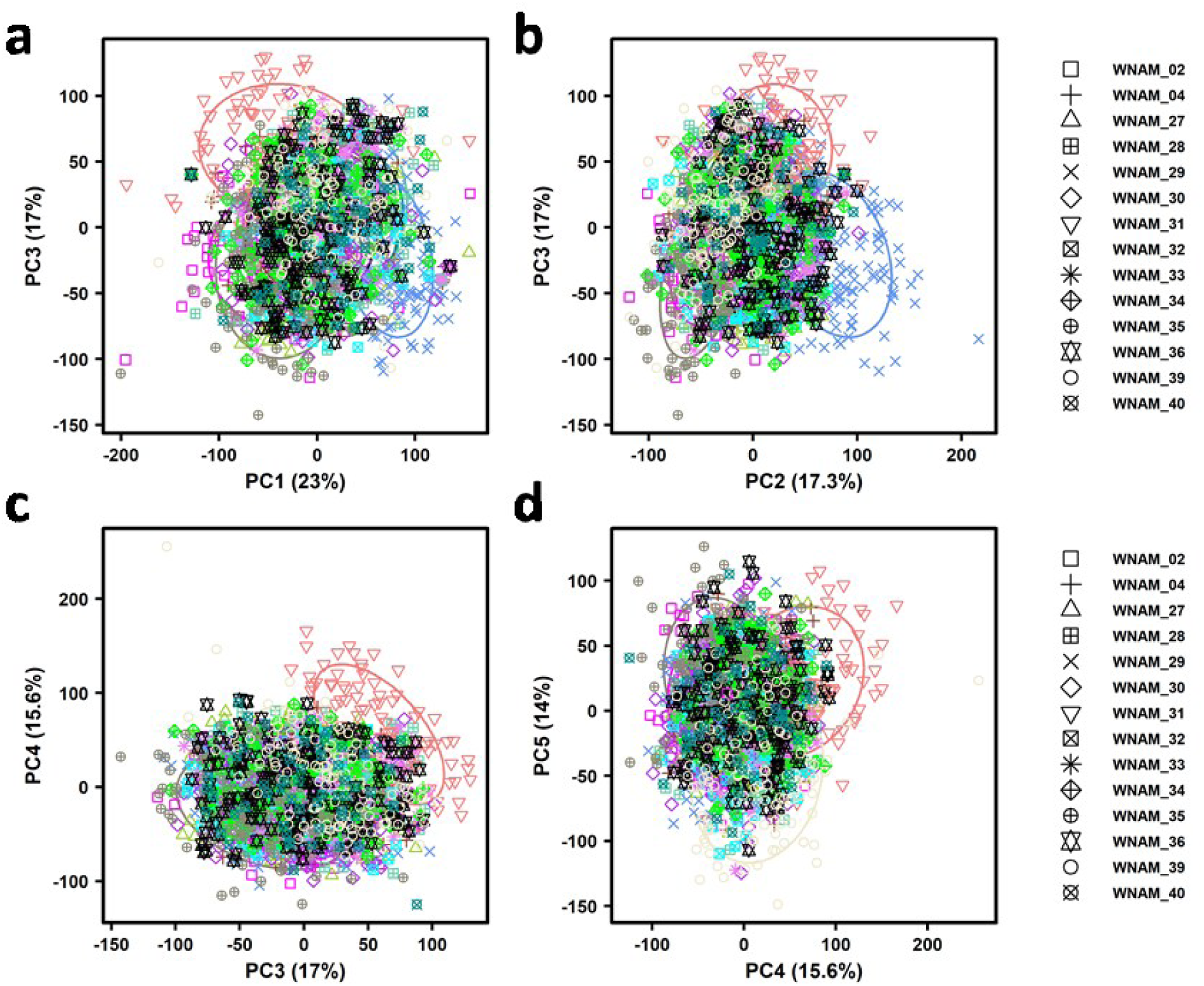
Higher-order principal component structure of the aus-NAM population. (a) PC1 (23.0%) vs PC3 (17.0%); (b) PC2 (17.3%) vs PC3 (17.0%); (c) PC3 (17.0%) vs PC4 (15.6%); and (d) PC4 (15.6%) vs PC5 (14.0%). Points represent individual RILs colored by family, while shapes correspond to NAM families as indicated in the legend on the right; ellipses denote group dispersion.

**Supplemental Fig. 4.**
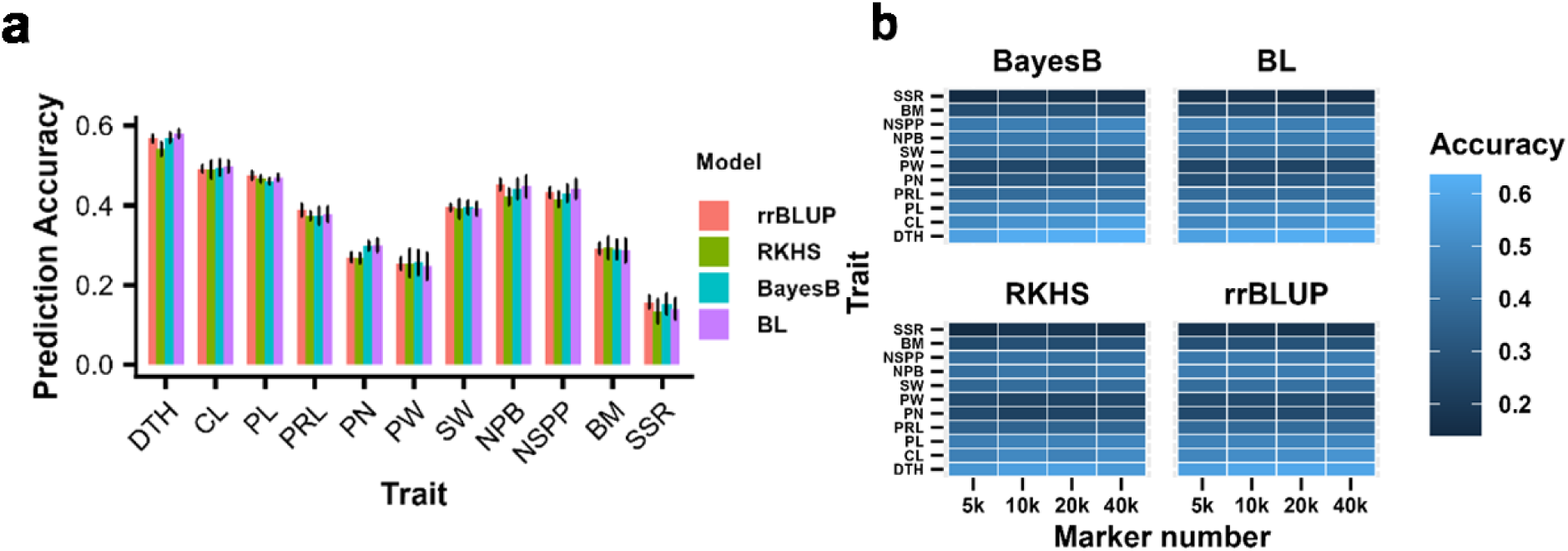
Prediction accuracy in aus-NAM. (a) Distribution of prediction accuracy across models and traits (mean ± SD). (b) Heatmap of prediction accuracy (trait × marker density × model)

**Supplemental Fig. 5.**
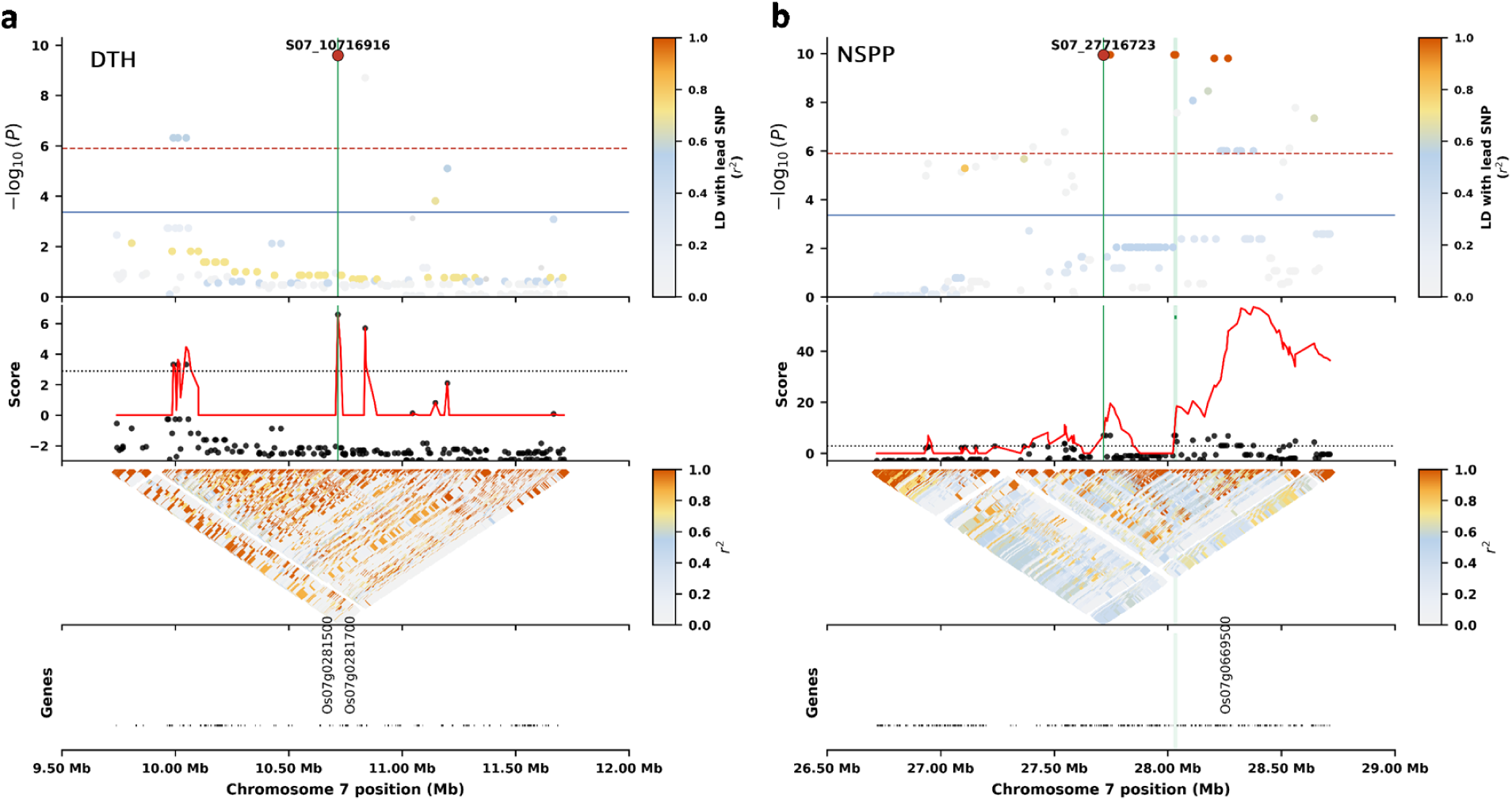
GWAS for agronomic traits. Local GWAS across genomic position with top panels showing SNP significance (−log₁₀(p); Manhattan plots) colored by linkage disequilibrium (LD; r²) relative to the lead SNP, middle panels showing the local score profile highlighting accumulation of association signal with the candidate interval indicated by a shaded region, and bottom panels showing pairwise LD (r²) as triangular heatmaps, revealing local haplotype structure and refinement of association peaks. (a) chromosome 7 region around S07_10716916 for days to heading (DTH), encompassing candidate genes including *OsAAO3* (Os07g0282300); (b) chromosome 7 region for number of spikelets per panicle (NSPP), encompassing candidate genes including *FRIZZY PANICLE* (*FZP*; Os07g0669500).

